# Complete Coding Genome Sequence for a Novel Multicomponent Kindia Tick Virus Detected from Ticks Collected in Guinea

**DOI:** 10.1101/2020.04.11.036723

**Authors:** Vladimir A. Ternovoi, Elena V. Protopopova, Alexander N. Shvalov, Mikhail Yu. Kartashov, Roman B. Bayandin, Tatyana V. Tregubchak, Sergey A. Yakovlev, Konstantin A. Nikiforov, Svetlana N. Konovalova, Valery B. Loktev, Alexander P. Agafonov, Rinat A. Maksyutov, Anna Yu. Popova

**Affiliations:** State Research Center for Virology and Biotechnology “Vector”, Koltsovo, Novosibirsk Region, Russia; Federal Government Health Institution Russian Research Anti-Plaque Institute “Microbe”, Saratov, Russia; Russian Federal Service for Surveillance on Consumer Rights Protection and Human Wellbeing, Moscow, Russia

## Abstract

Kindia tick virus (KITV) is a novel multicomponent virus first detected by direct sequencing of *Rhipicephalus geigyi* ticks in Guinea in 2017. Here, we present a complete coding genome sequence for all four segments of KITV/2017/1. This virus appears to be evolutionarily related to unclassified flaviviruses, such as Alongshan virus.

## ANNOUNCEMENT

The multicomponent (segmented) tick flaviviruses evolutionarily related to the unsegmented viruses of the genus *Flavivirus* have been recognized since only 2014 (1-4). This group of unclassified segmented tick-borne flaviviruses now includes Alongshan virus, Jingmen tick virus (JMTV) and Mogiana tick virus (MGTV), which have been detected in Asia, Europe and South America, respectively. These diverse and globally distributed viruses are capable of infecting a wide range of hosts, such as ticks, animals and humans (1–8). However, only a few complete genome sequences have been reported for JMTV (Kosovo and China), Alongshan virus (China) and MGTV (Brasilia).

Here, we report the complete coding genome sequences for the first African isolate, named Kindia tick virus (KITV), from *Rhipicephalus geigyi* collected from domestic cattle (*Bos taurus*) in Kindia, Guinea, West Africa. Twenty-six pools of 5 ticks each were frozen in liquid nitrogen, crushed by plastic pestles, homogenized in phosphate-buffered saline and used for RNA isolation with TRIzol reagent (Invitrogen Co., USA). Total RNA was quantified with a Qubit RNA Assay Kit (Invitrogen Co., USA) following the manufacturer’s instructions. RNA-seq libraries were constructed with an NEBNext Ultra RNA Library Prep Kit for Illumina (New England Biolabs). Sequencing was performed using a MiSeq Reagent Kit v3 for 600 cycles. Cutadapt (version 1.18) and SAMtools (version 0.1.18) were used to remove the Illumina adaptors and duplicate reads. After removing adapters, the read length was 108-118 bases, with the numbers of reads per pool ranging from 151,282 to 540,929. The contigs were assembled *de novo* using the MIRA assembler with default parameters (version 4.9.6). In five pools, we found Mogiana-like fragments using BLASTN. These fragments were aligned to the reference genome segments of Mogiana tick virus isolate MGTV/V4/11 (4). The average coverage of the four segments was 35, 18, 52, and 30, respectively. The complete sequences for KITV/2017/1 were verified by Sanger sequencing of Seg 2 (2 short fragments), Seg 3 (1 sf) and Seg 4 (2 sf), with gaps or low coverage, as previously described (9). Total RNA from tick pools positive for KITV/2017/1 was used for first-strand DNA synthesis using a Reverta-L Kit (Interlabservice, Russia). The KITV/2017/1 genetic material was amplified by PCR with specific primers designed based on draft NGS sequences (available on request), with subsequent PCR fragment isolation and sequencing. The sequences for two detected KITVs (KITV/2017/1 and KITV/2017/2) were also deposited in GenBank. The nucleotide identity between KITV/2017/1 and KITV/2017/2 was 99.7% (Seg 1), 99.4% (Seg 2), 98.1% (Seg 3), and 99.3% (Seg 4). KITV contains putative open reading frames (ORFs) congruent with JMTV and MGTV, namely, nonstructural protein 1 (Seg 1), VP1 (Seg 2), NSP2 (Seg 3), VP2 and VP3 (Seg 4). The sizes of the sequenced segments (ORFs) are 2968 (2743), 2805 (2262), 2667 (2427) and 2725 (2351) bases, with CG contents of 52.3%, 54.9%, 54.4% and 54.2% for each segment, respectively. The lengths of the 5’ UTR and 3’ UTR were different for each segment and were 97-156 and 121-387 bases, respectively. The 5’ UTR conservative motif GCAAGTGCA typical for JMTV was found in four segments, and the 3’ UTR conservative motifs GGCAAGTGC and CAAGTG were also found in Seg 2 and Seg 4 of KITV. The divergence between KITV and MGTV/JMTV based on nucleotide sequence was 7–28% (Seg 1), 7.7–20.6% (Seg 2), 2.3–29.3% (Seg 3), and 0.4–22% (Seg 4), and based on amino acid sequences, it was 3.1–21.8% (Seg 1), 3.4–20.3% (Seg 2), 1.3–20.8% (Seg 3), and 0.2–15% (Seg 4), as evaluated by BLAST. Given these similarities, we suggest that KITV is a new member of the multicomponent (segmented) tick-borne flavivirus group and possibly represents a new species together with MGTV.

### Data availability

GenBank accession numbers for the viral sequences: <u>MH678723</u> and <u>MH678727</u> (Seg 1), <u>MH678724</u> and <u>MH678728</u> (Seg 2), <u>MH678725</u> and <u>MH678729</u> (Seg 3), and <u>MH678726</u> and <u>MH678730</u> (Seg 4); complete sequences <u>MK673133</u> (Seg 1), <u>MK673134</u> (Seg 2), <u>MK673135</u> (Seg 3), and <u>MK673136</u> (Seg 4); and Sequence Read Archive accessions <u>SRX5930668</u> to <u>SRX5930671</u> under project <u>PRJNA545394</u>. The annotations have also been deposited into GenBank for these sequences.

## ACKNOWLEDGMENTS

The authors would like to gratefully acknowledge support from the Russian Federal Service for Surveillance on Consumer Rights Protection and Human Wellbeing, Project RP?2904, from 12/22/2017.

## REFERENCES

1. Qin XC, Shi M, Tian JH, Lin XD, Gao DY, He JR, Wang JB, Li CX, Kang YJ, Yu B, Zhou DJ, Xu J, Plyusnin A, Holmes EC, Zhang YZ. 2014. A tick-borne segmented RNA virus contains genome segments derived from unsegmented viral ancestors. Proc Natl Acad Sci U S A 111:6744–6749. https://doi.org/10.1073/pnas.1324194111.

2. Shi M, Lin XD, Vasilakis N, Tian JH, Li CX, Chen LJ, Eastwood G, Diao XN, Chen MH, Chen X, Qin XC, Widen SG, Wood TG, Tesh RB, Xu J, Holmes EC, Zhang YZ. 2015. Divergent viruses discovered in arthropods and vertebrates revise the evolutionary history of the *Flaviviridae* and related viruses. J Virol 90:659–669. https://doi.org/10.1128/JVI.02036-15.

3. Ladner JT, Wiley MR, Beitze lB, Auguste AJ, Dupuis AP, J., Lindquist ME, Sibley SD, Kota KP, Fetterer D, Eastwood G, Kimmel D, Prieto K, Guzman H, Aliota MT, Reyes D, Brueggemann EE, StJohn L, Hyeroba D, Lauck M,Friedrich TC, O’Connor DH, Gestole MC, Cazares LH, Popov VL, Castro-Llanos F, Kochel TJ, Kenny T, White B, Ward MD, Loaiza JR, Goldberg TL,Weaver SC, Kramer LD, Tesh RB, Palacios G. 2016. A multicomponent animal virus isolated from mosquitoes. Cell Host Microbe 20:357–367. https://doi.org/10.1016/j.chom.2016.07.011.

4. Villa EC, Maruyama SR, de Miranda-Santos IKF, Palacios G, Ladner JT. 2017. Complete Coding Genome Sequence for Mogiana Tick Virus, a Jingmenvirus Isolated from Ticks in Brazil. Genome Announc 5(18). pii: e00232–17. https://doi.org/10.1128/genomeA.00232-17.

5. Shi M, Lin XD, Vasilakis N, Tian JH, Li CX, Chen LJ, Eastwood G, Diao XN, Chen MH, Chen X, Qin XC, Widen SG, Wood TG, Tesh RB, Xu J, Holmes EC, Zhang YZ. 2015. Divergent viruses discovered in arthropods and vertebrates revise the evolutionary history of the *Flaviviridae* and related viruses. J Virol 90:659–669. https://doi.org/10.1128/JVI.02036-15

6. Meng F, Ding M, Tan Z, Zhao Z, Xu L, Wu J, He B, Tu C. 2019. Virome analysis of tick-borne viruses in Heilongjiang Province, China. Ticks Tick Borne Dis 10(2):412–420. https://doi.org/10.1016/j.ttbdis.2018.12.002.

7. Emmerich P, Jakupi X, von Possel R, Berisha L, Halili B, Günther S, Cadar D, Ahmeti S, Schmidt-Chanasit J. 2018. Viral metagenomics, genetic and evolutionary characteristics of Crimean-Congo hemorrhagic fever orthonairovirus in humans, Kosovo. Infect Genet Evol 65:6–11. https://doi.org/10.1016/j.meegid.2018.07.010.

8. Wang ZD, Wang B, Wei F, Han SZ, Zhang L, Yang ZT, Yan Y, Lv XL, Li L, Wang SC, Song MX, Zhang HJ, Huang SJ, Chen J, Huang FQ, Li S, Liu HH, Hong J, Jin YL, Wang W, Zhou JY, Liu Q.A. 2019. New Segmented Virus Associated with Human Febrile Illness in China. N Engl J Med 380(22):2116–2125. https://doi.org/10.1056/NEJMoa1805068.

9. Ponomareva EP, Ternovoi VA, Mikryukova TP, Protopopova EV, Gladysheva AV, Shvalov AN, Konovalova SN, Chausov EV, Loktev VB. 2017. Adaptation of tick-borne encephalitis virus from human brain to different cell cultures induces multiple genomic substitutions. Arch Virol 162(10):3151–3156. https://doi.org/10.1007/s00705-017-3442-x

